# Synaptotagmin 2 is ectopically overexpressed in excitatory presynapses of a widely used CaMKIIα-Cre mouse line

**DOI:** 10.1101/2021.06.30.450492

**Authors:** Ken Matsuura, Haytham Mohamed Aly Mohamed, Mohieldin Magdy Mahmoud Youssef, Tadashi Yamamoto

## Abstract

The CaMKIIα-Cre mouse line, one of the earliest established Cre driver lines, has resulted in over 800 papers to date. Here, we demonstrate that the second most widely used CaMKIIα-Cre line, Tg(Camk2a-cre)2Gsc (or CamiCre), shows ectopic overexpression of synaptotagmin 2, the most efficient Ca^2+^ sensor for fast synchronous neurotransmitter release, in excitatory presynapses of Cre^+^ brains. Moreover, RNA-seq analysis showed aberrant expression in Cre^+^ hippocampus, including upregulation of immediate early genes, such as *Arc* and *Fos*, and genes presumably derived from bacterial artificial chromosome transgene, such as *Slc6a7*. Most importantly, CamiCre^+^ mice showed functional phenotypes, such as hyperactivity and enhanced associative learning, suggesting neural activities are affected. These unexpected results suggest difficulties in interpreting results from studies using the CamiCre line and raise awareness of potential pitfalls in the use of Cre driver lines in general.

## Introduction

The Cre-loxP system is one of the most powerful and commonly used techniques in neurogenetics, as well as in broad biomedical studies, ranging from spatial and/or temporal gene-knockout (KO)s/knockin (KI)s, neural pathway tracing, to optogenetics (Branda and Dymecki, 2004; Daigle et al., 2018; Gerfen et al., 2013). The CaMKIIα-Cre mouse was the first Cre driver line in neurogenetics that showed successful recombination of DNA between two palindromic loxP sites inserted in critical positions of a gene of interest by the bacteriophage DNA-recombinase Cre to generate conditional gene-KO in principal neurons of mouse forebrain (Tsien et al., 1996a; Tsien et al., 1996b). There are currently 37 different CaMKIIα-Cre lines available, with 31 of them used in one or more papers (Fig S1A) according to the Mouse Genome Informatics (MGI) database (Bult et al., 2019) (http://www.informatics.jax.org/searchtool/Search.do?query=Camk2a-cre&page=featureList). These mouse lines are used in various studies addressing gene functions in electrophysiology, learning and memory, circadian rhythm, metabolism regulation, neuropsychiatric disease models, and so forth, in over 800 publications to date (Fig. S1A).

CamiCre is the second most widely used line next to the original CaMKIIα-Cre line, Tg(Camk2a-cre)T29-1Stl (T29-1) (Tsien et al., 1996a; Tsien et al., 1996b), and has been used in at least 126 papers since its first appearance in 2001 (Casanova et al., 2001) (Fig. S1A), many of them in high impact journals (http://www.informatics.jax.org/reference/allele/MGI:2181426?typeFilter=Literature#myDataTable=results%3D25%26startIndex%3D0%26sort%3Dyear%26dir%3Ddesc%26typeFilter%3DLiterature). The main feature of CamiCre mice is the incorporation of bacterial artificial chromosome (BAC) technology, which improves fidelity of expression of Cre recombinase under control of CaMKIIα regulatory sequences, since BACs contain most if not all required regulatory elements (Casanova et al., 2001; Gong et al., 2007; Heintz, 2001). Another feature is the use of improved Cre (iCre) to maximize Cre expression and activity (Branda and Dymecki, 2004; Casanova et al., 2001; Shimshek et al., 2002). Expression of Cre is mainly restricted to excitatory principal neurons in the forebrain, but is also enriched in the suprachiasmatic nucleus in the hypothalamus, which makes it a valuable tool for research in circadian rhythm and/or regulation of metabolism (Cedernaes et al., 2019; Godinho-Silva et al., 2019).

Synaptotagmin 2 (Syt2) is one of three synaptotagmins (Syt1, 2 and 9) that act as Ca^2+^ sensors for fast synchronous neurotransmitter release from presynaptic terminals. Ca^2+^ binding to Syt1, 2 or 9 induces their interaction with the soluble NSF attachment protein receptor (SNARE) complexes, thereby displacing complexins from assembled SNARE complexes to trigger neurotransmitter release (Chen et al., 2017; Xu et al., 2007). These three synaptotagmins form a hierarchy of Ca^2+^ sensors with distinct properties, with Syt2 being the fastest isoform (30% faster than Syt1), and Syt9, the slowest (Xu et al., 2007). Syt2 also triggers release with shorter latency and higher temporal precision and mediates faster vesicle pool replenishment than Syt1 (Chen et al., 2017). In mouse forebrain, Syt1 and Syt2 have complementary expression patterns and are localized predominantly to different subsets of synapses (Fox and Sanes, 2007). Syt2 is selectively expressed in inhibitory neurons in this region and preferentially in large axosomatic synapses (Fox and Sanes, 2007; Pang et al., 2006). It is postulated that these differential expression patterns warrant firing properties required for different neuronal types, e.g., Syt2 is the only isoform expressed in the calyx synapse, which relies on very fast signaling (Xu et al., 2007).

Here, we show that CamiCre^+^ mice show ectopic overexpression of Syt2 in excitatory presynapses in the forebrain and functional phenotypes, such as hyperactivity and enhanced learning, which may compromise research using CamiCre mice. These unexpected and problematic phenotypes in CamiCre mice shed light on the importance of better phenotypic characterization of Cre driver lines and the importance of adopting proper controls, including the incorporation of Cre^+^ controls in experimental designs, which are often neglected.

## Results and Discussion

### Syt2 is significantly upregulated in excitatory presynapses of CamiCre^+^ forebrain

In the course of a project utilizing CamiCre mice to generate conditional KO mice for the gene of interest, we noticed upregulation of Syt2 in both transcriptomic and proteomic analyses of KO brain compared to *Gene*^*flox/flox*^; CamiCre^−^ controls. We were initially excited, but as a result of subsequent qPCR and Western blot experiments that included *Gene*^+/+^; CamiCre^+^ controls, we soon realized that this upregulation was independent of our gene of interest. Therefore, we deemed it would be important for our project and possibly to any research involving CamiCre mice to characterize this Cre driver line in more detail first.

Using the original CamiCre line obtained from the European Mouse Mutant Archive (EMMA), which was backcrossed seven times onto a C57BL/6J genetic background, qPCR results confirmed that *Syt2* mRNA was significantly upregulated more than 5-fold in CamiCre^+^ hippocampus compared to wild-type (WT) littermate (CamiCre^−^) controls (Fig. 1A). This upregulation was likely due to increased transcriptional activity suggested by significant upregulation of *Syt2* premRNA (Fig 1A). This upregulation was also confirmed at the protein level by Western blotting of lysates from different brain regions (Fig. 1B). Syt2 was overexpressed more than 7-fold in the hippocampus and about 3-fold in the olfactory bulb and cerebral cortex. However, no significant upregulation was observed in caudal brain regions, such as the midbrain, cerebellum, or medulla oblongata (Fig. 1C, D). Syt1 expression was not changed in any brain region analyzed (Fig. 1A and Fig. S2A, B). As the effect was restricted to the forebrain of CamiCre^+^ mice, mimicking the CaMKIIα promoter-driven Cre expression pattern, we wondered whether simple Cre expression could affect expression of Syt2. We analyzed two additional driver lines, the original CaMKIIα-Cre line T29-1 and Emx1^tm1(cre)Ito^ (Iwasato et al., 2000), which also express Cre in neurons of the forebrain. These mice did not show any upregulation of Syt2 in the hippocampus of Cre^+^ mice in either mRNA (Fig 1A) or protein (Fig 1E) levels. However, since the Cre expression level was much lower in these two lines compared to CamiCre (likely due to the iCre effect), we also adopted adeno-associated virus (AAV)-mediated Cre expression (AAV-hSyn-Cre) in hippocampal neurons of WT mice to see whether high expression of Cre would affect the Syt2 expression level (Fig 1E). The results suggested that simple expression of Cre in neurons, regardless of its amount, would not affect the Syt2 expression level. This was good news, as the results suggested that using the Cre-loxP system in neurons in general will not be compromised by the Syt2 overexpression.

**Figure 1.**
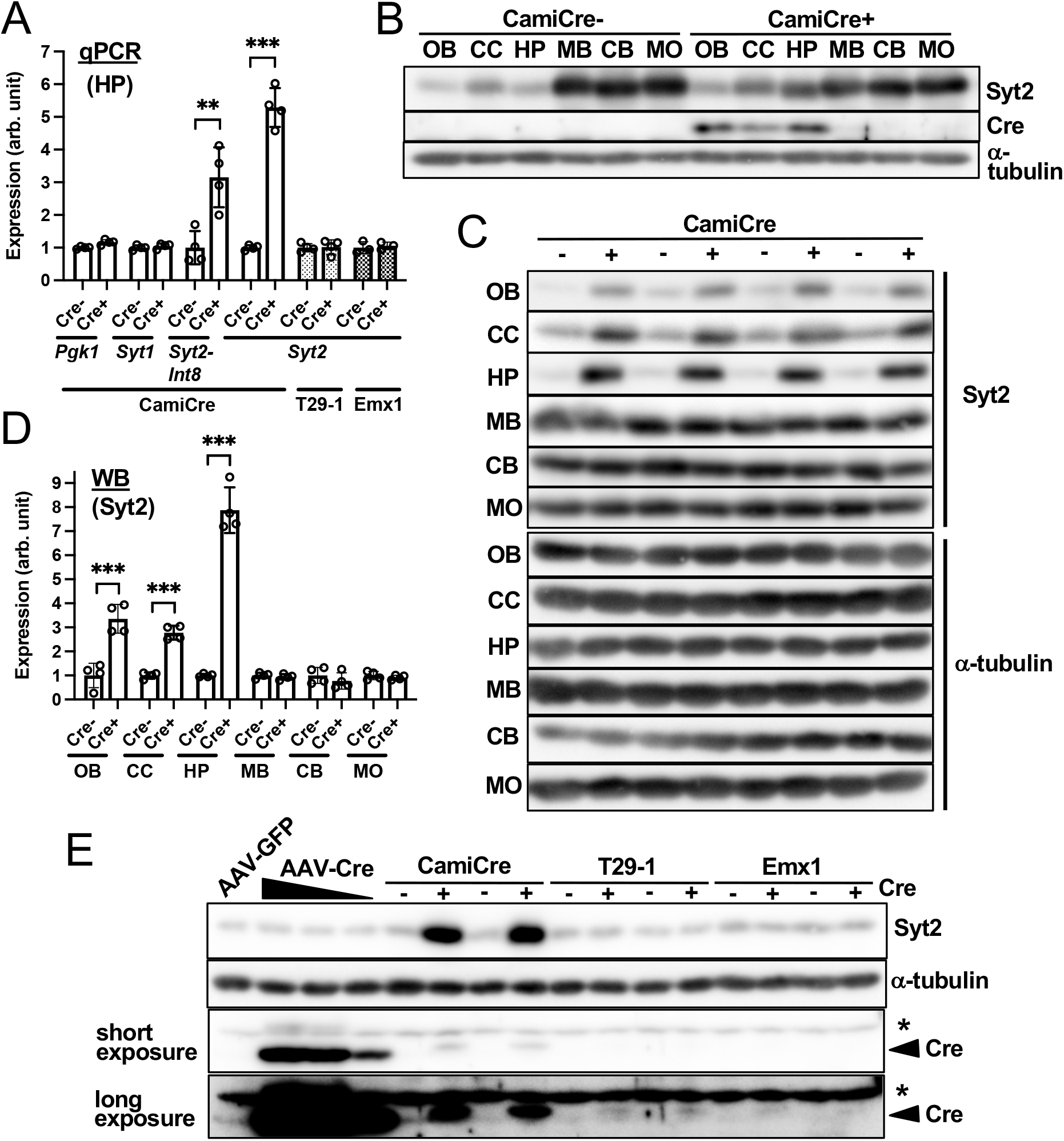
Syt2 is significantly upregulated in CamiCre^+^ forebrain. (**A**) qPCR analysis of mRNA expression in the hippocampus in indicated mouse lines. *Syt2-Int8* represents premRNA of *Syt2*. Values were normalized by *Gapdh* expression level and are shown as fold increases against Cre-controls. *Pgk1* is another housekeeping gene. (**B, C**) Western blotting images of different brain regions. CB, cerebellum; CC, cerebral cortex; HP, hippocampus; MB, midbrain; MO, medula oblongata; OB, olfactory bulb. (**D**) Quantification results of C. Values were normalized against α-tubulin expression level and are shown as fold increases against Cre-controls. (**E**) Western blotting images of hippocampal lysates showing Syt2 expression levels in various Cre-expressing methodologies. Viral concentration was diluted 2x in three steps for AAV-hSyn-Cre expression. *, nonspecific band. Mean ± SD is shown. **P < 0.01; ***P < 0.001. N = 4 (biological replicates). Statistical tests were performed using two-tailed, unpaired t-tests.

The next question was whether Syt2 overexpression occurs in its native location, the synaptic terminals of inhibitory neurons (Fox and Sanes, 2007; Pang et al., 2006), or in other cell types and/or subcellular structures. Immunostaining of Syt2 in the hippocampus of CamiCre^+^ mice showed a striking difference from WT controls (Fig.2A). Syt2 showed strong uniform staining in the neuropil area of the CamiCre^+^ hippocampus, compared to much more limited and sporadic staining in the neuropil area of WT controls. High resolution colocalization analysis using a Confined Displacement Algorithm (Ramirez et al., 2010) in the neuropil region of hippocampus CA1 and CA3 demonstrated significant correlation and colocalization of Syt2 with the vesicular glutamate transporter 1 (VGluT1), a glutamatergic presynapse marker (Fig. 2B, C). More than 80% of Syt2 signals colocalized with VGluT1 in the neuropil region of CamiCre^+^ hippocampus. Syt2 staining in axosomatic synapses surrounding pyramidal neurons was similar between CamiCre^+^ and WT hippocampi (Fig. 2B). These results suggested that Syt2 is primarily overexpressed ectopically in presynapses of excitatory neurons in CamiCre^+^ mice.

**Figure 2.**
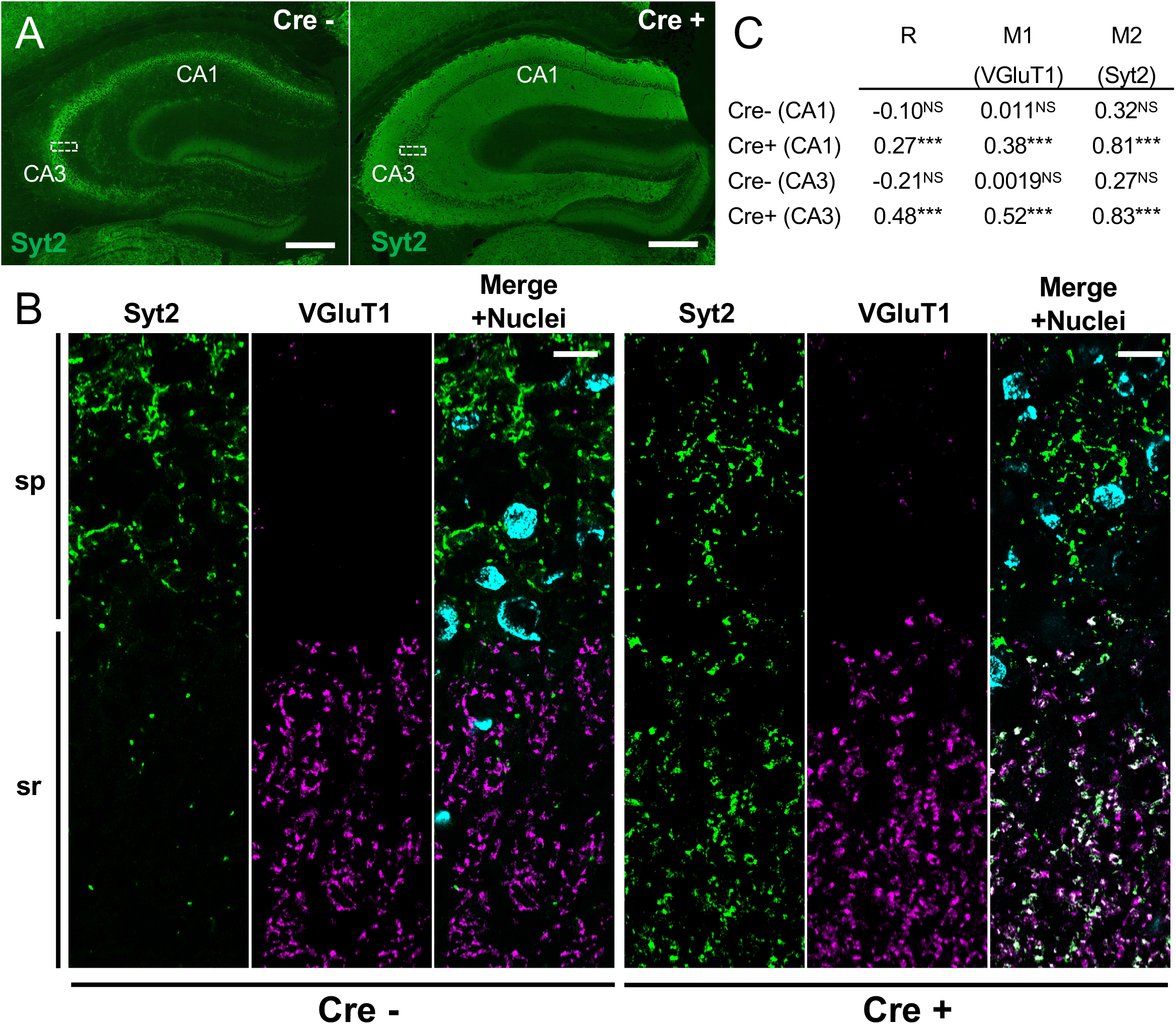
Syt2 is ectopically overexpressed in excitatory presynapses of CamiCre^+^ hippocampi. (**A**) Immunofluorescent images of Syt2 staining in hippocampi of CamiCre mice. (**B**) Super-resolution images of the hippocampal CA3 region corresponding to the approximate areas indicated by the white dotted boxes in A are shown. sp, stratum pyramidale; sr, stratum radiatum. (**C**) Colocalization was analyzed using the Confined Displacement Algorithm on the neuropil area of CA1 and CA3. R, Pearson correlation coefficient; M1 and M2, Manders coefficient. NS, Not significant; ***P < 0.001 (the significance of correlation or colocalization were calculated by comparing original images to random displacement images). Scale bars: (A) 250 μm; (B) 10 μm.

### CamiCre^+^ mice show aberrant gene expression, hyperactivity, and enhanced learning in fear conditioning

We further asked whether there are additional differentially expressed genes in CamiCre^+^ hippocampus using RNA-seq analysis. In addition, we performed parallel analysis on the most widely used T29-1 line to examine whether there are any overlaps between the two CaMKIIα-Cre lines. Differential expression (DE) analysis suggested upregulation of 329 genes in CamiCre^+^ hippocampus and 247 in T29-1 Cre^+^ hippocampus. Moreover, 172 and 226 genes were downregulated in CamiCre^+^ and T29-1 Cre^+^ hippocampus, respectively (Table S1 and S2). However, there were no significant overlaps of differentially expressed genes between the two lines (Fig. S3A).

Analysis of CamiCre^+^ hippocampus indicated that 4 genes, including *Syt2*, were overexpressed with remarkably high confidence (Fig. 3A and Table S1). Three others were *Slc6a7*, *Arsi*, and *Cdx1*. We found that these 3 genes were located in the genomic region within the arms of the CaMKIIα-iCre BAC transgene used to generate CamiCre mice (Casanova et al., 2001) (Fig.3 B). Thus, overexpression of these 3 genes in CamiCre^+^ mice is presumably derived from the BAC transgene. This issue may be common to Tg(Camk2a-cre/ERT2)2Gsc, the most widely used inducible type of CaMKIIα-Cre line (Fig. S1A), as it uses a common BAC clone structure (Erdmann et al., 2007). Although the mechanism of *Syt2* overexpression is not clear, one plausible explanation is that the BAC transgene is inserted in the promoter region of *Syt2* to disrupt its native suppression machinery and possibly its expression controlled by regulatory elements of *CaMKIIα*, *Slc6a7*, and/or *Cdx1*. These 3 genes have mostly forebrain-specific expression, according to the Allen Mouse Brain Atlas (http://mouse.brain-map.org). This model accords well with the observed overexpression of Syt2 in the forebrain (Fig. 1B-D). Alternatively, it could be related to the unintended recombination through pseudo-loxP sites, which is postulated to cause phenotypes for Nestin-Cre (Harno et al., 2013) and Lck-Cre mice (Carow et al., 2016) in Cre high-expressing cells. However, at least Cre overexpression in hippocampal neurons by AAV infection did not result in change in Syt2 expression level (Fig. 1E). In the T29-1 Cre^+^ hippocampus, *St6gal2*, a gene encoding beta-galactoside alpha-2,6-sialyltransferase 2, was overexpressed (3.6-fold of controls) with very high confidence (Table S2). The physiological role of *St6gal2* is not well studied, but it is implicated in schizophrenia (Ikeda et al., 2010) and autism (Guo et al., 2017). Mechanism for this upregulation is also not clear, but likely related to the transgene insertion effect, as low Cre expression in T29-1 is unfavorable for recombination of pseudo-loxP sites, and *St6gal2* upregulation is not observed in CamiCre^+^ mice.

**Figure 3.**
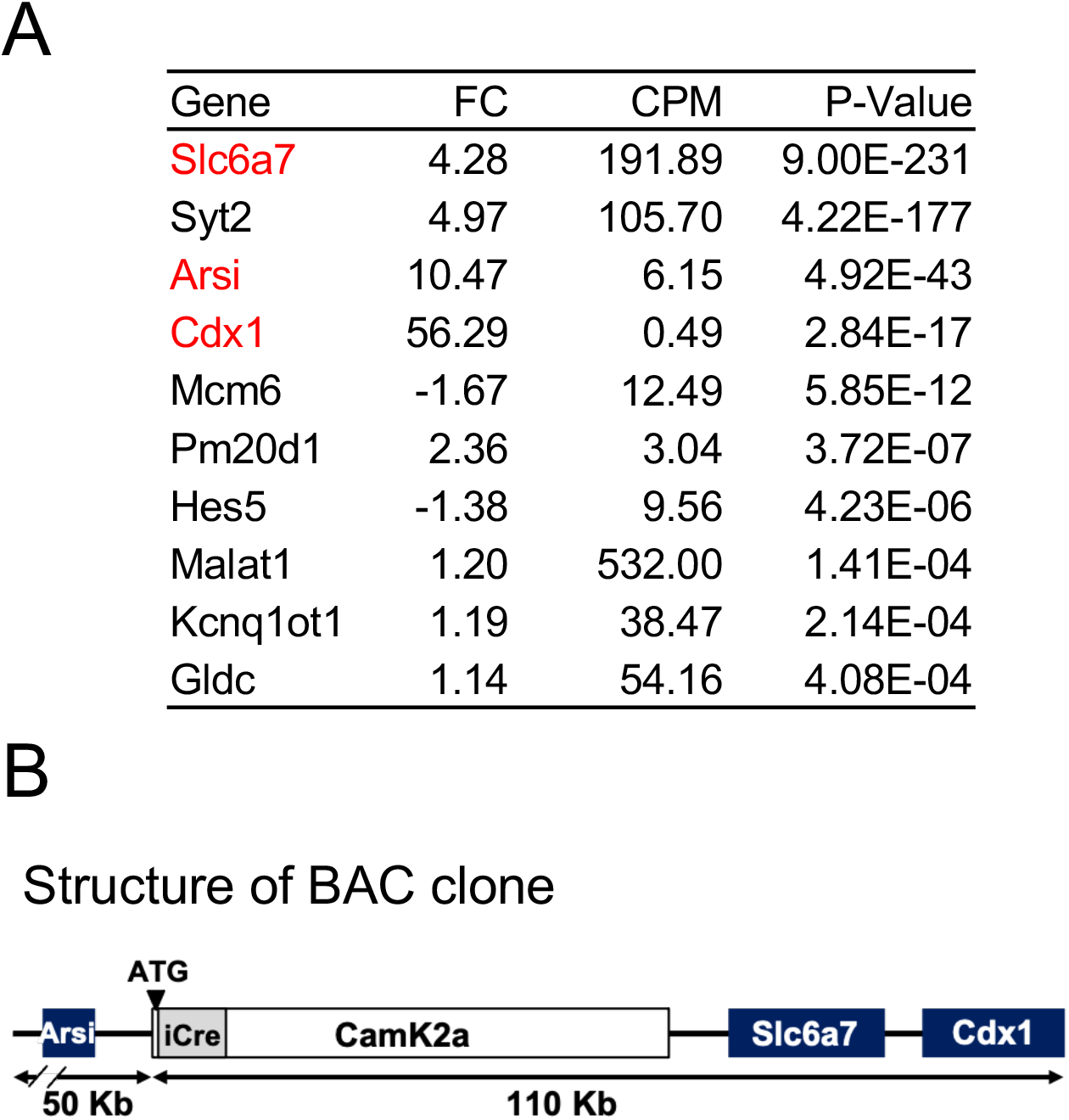
Aberrant gene expression in CamiCre^+^ hippocampi. (**A**) The top 10 differentially expressed genes in CamiCre^+^ hippocampi, ranked by P-values of exact tests in DE analysis of RNA-seq data. N = 4. FC, fold changes of expression in CamiCre^+^ mice compared to WT littermates; CPM, average counts per million of all samples. Genes located on the BAC transgene of CamiCre mice are indicated in red. (**B**) Schematic representation of the BAC clone used to generate CamiCre mice.

We performed Ingenuity Pathway Analysis (IPA) on candidate differentially expressed genes to see whether there is a specific group of genes in signaling pathways that is consistently up- or downregulated. The results suggested upregulation of the CREB signaling pathway in CamiCre^+^ hippocampus and we noticed that most of the top-ranking pathways identified included upregulation of immediate early genes (IEGs) such as *Arc*, *Fos*, *Egr2*, and *Nr4a1* (Table S3). IEGs are rapidly and transiently transcribed genes in response to neuronal stimulation without a requirement for new protein synthesis and are thought to reflect neuronal activity (Yap and Greenberg, 2018). The extent of upregulation of IEGs in CamiCre^+^ hippocampus was relatively small (20-70% upregulated) in our bulk analysis, but the tendency was clear compared to results from T29-1 Cre^+^ hippocampus (Fig. S3B and Tables S1 and S2) and the upregulation can be confirmed by qPCR analysis (Fig S3C). These results suggest that neuronal networks in CamiCre^+^ hippocampus tend to be more activated than in their WT littermates, which is consistent with the idea that the faster release kinetics of overexpressed Syt2 result in more efficient neuronal activity. In addition, *Slc6a7*, which functions as an L-proline transporter, may also be involved, as it is postulated to positively regulate glutamatergic neurotransmission (Schulz et al., 2018). In the IPA upstream analysis of the T29-1 Cre^+^ hippocampus, estrogen receptor 1 (ESR1) signaling appeared most likely to be affected (Table S4).

Finally, and most importantly, through a battery of common behavioral tests, we asked whether CamiCre^+^ mice show any functional phenotypes. CamiCre^+^ mice and their littermate controls were examined in open field, elevated plus maze, y-maze, light-dark transition, cued and contextual fear conditioning, and forced swim tests. CamiCre^+^ mice traveled significantly farther in the 15-min open field test (Fig. 4A). This was due to enhanced speed and extended duration per movement episode (Fig. 4B-D). CamiCre^+^ mice did not show a significant difference in the time spent near the center in the open field test, the time spent in the open arms in the elevated plus maze, or the time spent in the light box in a light-dark transition test (Fig. S4A-C). These results all suggest normal anxiety levels in CamiCre^+^ mice. They also did not show a significant difference in the forced swim test (Fig. S4D), which is used to test despair and depression. However, despite their hyperactivity and normal anxiety levels, CamiCre^+^ mice showed more freezing in both cued and contextual fear conditioning (Fig. 4E-H), which is indicative of better associative learning. On the other hand, they did not show significant differences in short-term spatial working memory in the y-maze test (Fig. S4E), but showed increased arm entry numbers, reflecting their hyperactivity (Fig. S4F). The hyperactivity and enhanced associative learning phenotypes may also be attributed to overexpression of Syt2 in excitatory presynapses, which may contribute to faster and more efficient neuronal signaling in CamiCre^+^ brains. Upregulation of *Slc6a7* may also be involved, as *Slc6a7*-deficient mice show reduced locomotor activity and impaired memory extinction (Schulz et al., 2018). *Slc6a7* is also implicated in autism, schizophrenia, and intellectual disability (Schulz et al., 2018).

**Figure 4.**
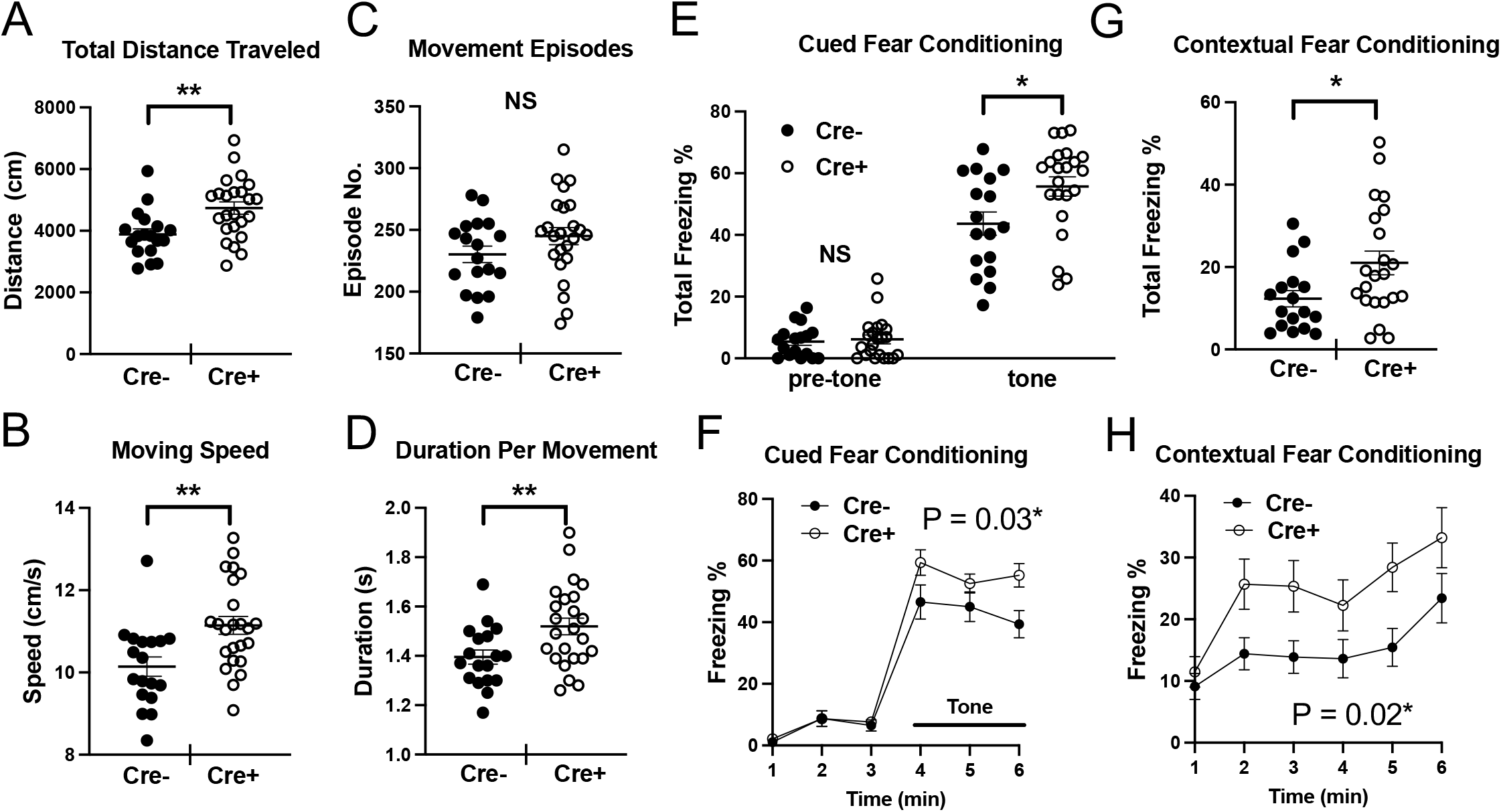
CamiCre^+^ mice show hyperactivity and enhanced learning in fear conditioning. (**A**-**D**) Open field test. (**E-H**) Cued and contextual fear conditioning. Mean ± SEM is shown. NS, not significant; *P < 0.05; **P < 0.01. P values labelled in the line graphs indicate the genotype effect of two-way ANOVA; (F) F (1, 37) = 5.023, (H) F (1, 37) = 6.204. N = 18 for Cre^−^, N = 24 for Cre^+^ for open field test and N = 17 for Cre^−^, N = 22 for Cre^+^ for cued and contextual fear conditioning. Mice that showed significant freezing before the first conditioned stimulus were excluded from the analysis.

### Potential problems in using CamiCre mice or Cre driver lines in general and ways to minimize them

In this report, we demonstrated that Syt2 is ectopically overexpressed in excitatory presynapses of CamiCre^+^ mice. Moreover, transcriptomic analyses indicate that there are many differentially expressed genes in CamiCre^+^ hippocampi, including the 3 genes located in the CaMKIIα-iCre BAC transgene. The biggest drawback is that activities of neural circuits of CamiCre^+^ mice are likely affected, as suggested by their hyperactivity and better learning in the fear conditioning test. These properties of CamiCre mice clearly pose a problem when utilizing them as Cre drivers, because the purpose is usually to examine the effect of the Cre-driven KO/KI of a gene of interest on neuronal function and its output behavior. Having said that, phenotypes of transcriptomic analyses and behavioral tests shown here could be considered mild compared to many severe phenotypes reported in the literature using CamiCre mice. Therefore, this work does not necessarily invalidate all previous reports using CamiCre mice. Nevertheless, phenotypes shown here are not trivial either, and results obtained using CamiCre mice should be interpreted with caution.

We have read at least two papers that report overexpression of Syt2 as a phenotype reflecting deletion of a gene of interest using the CaMKIIα-Cre line that do not clearly state which specific lines they used, but one shows a similar ectopic overexpression pattern in the hippocampus (Park et al., 2019) and the other shows concurrent upregulation of *Slc6a7* and *Arsi* (Wrackmeyer et al., 2019), suggesting use of the CamiCre line. As these two papers only used *Gene*^*flox/flox*^ controls, the results should be confirmed with Cre^+^ controls.

One of the simplest ways to minimize problems inherited in a Cre driver line is to include Cre^+^ controls. Various authors have urged routine use of Cre^+^ controls to circumvent numerous pitfalls in the Cre-loxP system and to correctly interpret data from conditional knockout studies (Becher et al., 2018; Carow et al., 2016; Harno et al., 2013; Lee et al., 2006; Schmidt-Supprian and Rajewsky, 2007). Unfortunately, however, these recommendations are still often neglected. In the case of CamiCre mice, only 15 out of 126 papers clearly mention use of Cre^+^ controls (Fig. S1B) and *Gene*^*flox/flox*^ or *Gene*^*flox*/+^; Cre^−^ controls are most frequently used.

Although Cre^+^ controls could clarify whether Cre^+^ itself affects phenotypes of conditional mutant mice to a certain extent, there is a limit. It is possible that phenotypes inherited in Cre driver lines could enhance or cancel true phenotypes of mutations of interest. Moreover, synergistic phenotypes of Cre mice with mutated genes of interest, which may happen in unpredictable mechanisms, could be manifested. These results might not always be clarified by Cre^+^ controls. One way to circumvent this issue is to find Cre driver lines that are established to have minimal phenotypes in systems of interest. However, using Cre^+^ controls is still important, as phenotypes could change with age, genetic background, housing environment, and experimenter handling of the mice. In cases in which researchers need to use particular Cre driver line with a potentially problematic phenotype for a specific expression region and/or Cre recombination efficiency, another way to circumvent the issue is to show similar results in different systems, such as: different Cre line(s) with the same or different promotors, conventional KO lines (if applicable), AAV-Cre-mediated KO mice, or RNAi/CRISPR-Cas9 mediated KDs/KOs in cell cultures (if applicable). These additional data could give assurance that results are independent of properties of a problematic Cre driver line.

To our knowledge, while Cre driver lines are well characterized for Cre expression patterns, many are not well characterized for phenotypes unless they are obvious, and most of genomic locations of transgene insertion are not known. This work shows the effectiveness of utilizing comprehensive transcriptome analysis on a Cre driver line for screening extreme changes in gene expression, which could result from transgene insertions, unintended recombination at pseudo-loxP sites, or even Cre toxicity (Becher et al., 2018; Carow et al., 2016; Harno et al., 2013; Lee et al., 2006; Schmidt-Supprian and Rajewsky, 2007). General assessment of the status of a tissue or a cell type of interest by pathway analyses is also informative. Perhaps creating a database of DE analyses for Cre driver lines would benefit researchers in choosing the right driver line for their research and the appropriate interpretation of their results. We hope this work will raise awareness of potential caveats in the Cre-loxP system in general, and will encourage researchers to be cautious in designing control experiments to maximize the usefulness of the system and validity of their results.

## Acknowledgements

We thank Drs. Shigeyoshi Itohara and Takuji Iwasato (RIKEN Brain Science Institute) for providing Emx1^tm1(cre)Ito^ mice. We thank Dr. Bernd Kuhn (Optical Neuroimaging Unit, OIST) for technical support on AAV injections. pENN.AAV.hSyn.Cre.WPRE.hGH and pAAV.hSyn.eGFP.WPRE.bGH were gifts from James M. Wilson. We are grateful for the help and support provided by the Animal Resource Section and the Sequencing Section of the Research Support Division at Okinawa Institute of Science and Technology Graduate University (OIST). This work was supported by JSPS KAKENHI Grant Numbers JP19K07394.

## Author contributions

Conceptualization: KM

Methodology: KM, HMAM, MMMY

Investigation: KM, HMAM, MMMY

Visualization: KM

Funding acquisition: KM, TY

Project administration: KM, TY

Supervision: KM, TY

Writing – original draft: KM

Writing – review & editing: KM, HMAM, MMMY, TY

## Data and materials availability

Most data generated or analyzed during this study are included in this article and its supplementary information files. The RNA-seq datasets used in the current study will be deposited in public database (GEO, NCBI). All materials are readily available from commercial sources or from our lab for reasonable requests.

---

**Figure S1.**
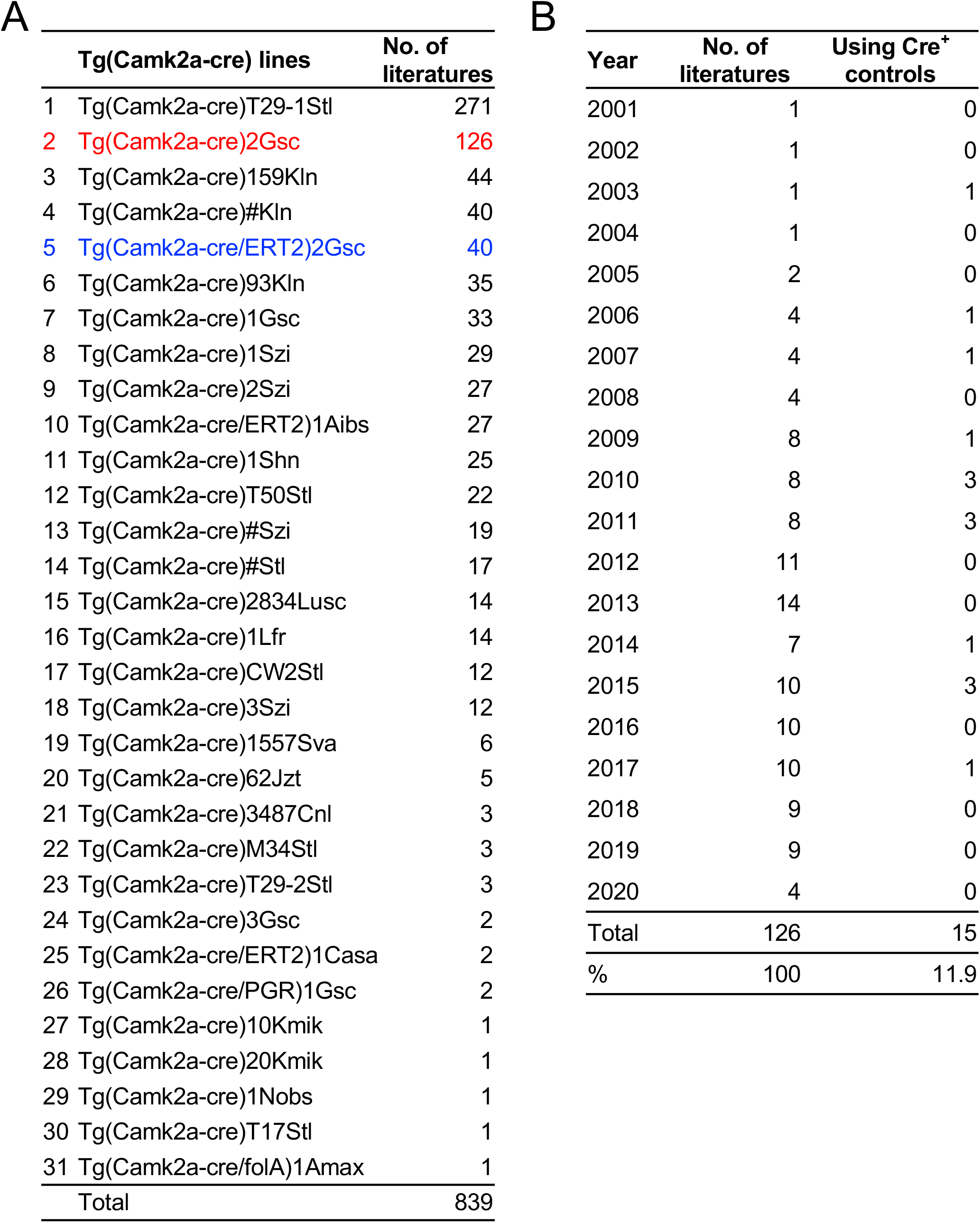
Literature survey. (**A**) Numbers of publications using specific Camk2a-Cre transgenic mouse lines from 1996 to 2020. Numbers are based on literature information from MGI. The CamiCre line is highlighted in red and the inducible Cre expression line that uses the same BAC clone strategy with the CamiCre line is highlighted in blue. (**B**) Numbers of publications stating the use of Cre-expressing controls for studies utilizing the CamiCre line in each publication year.

**Figure S2.**
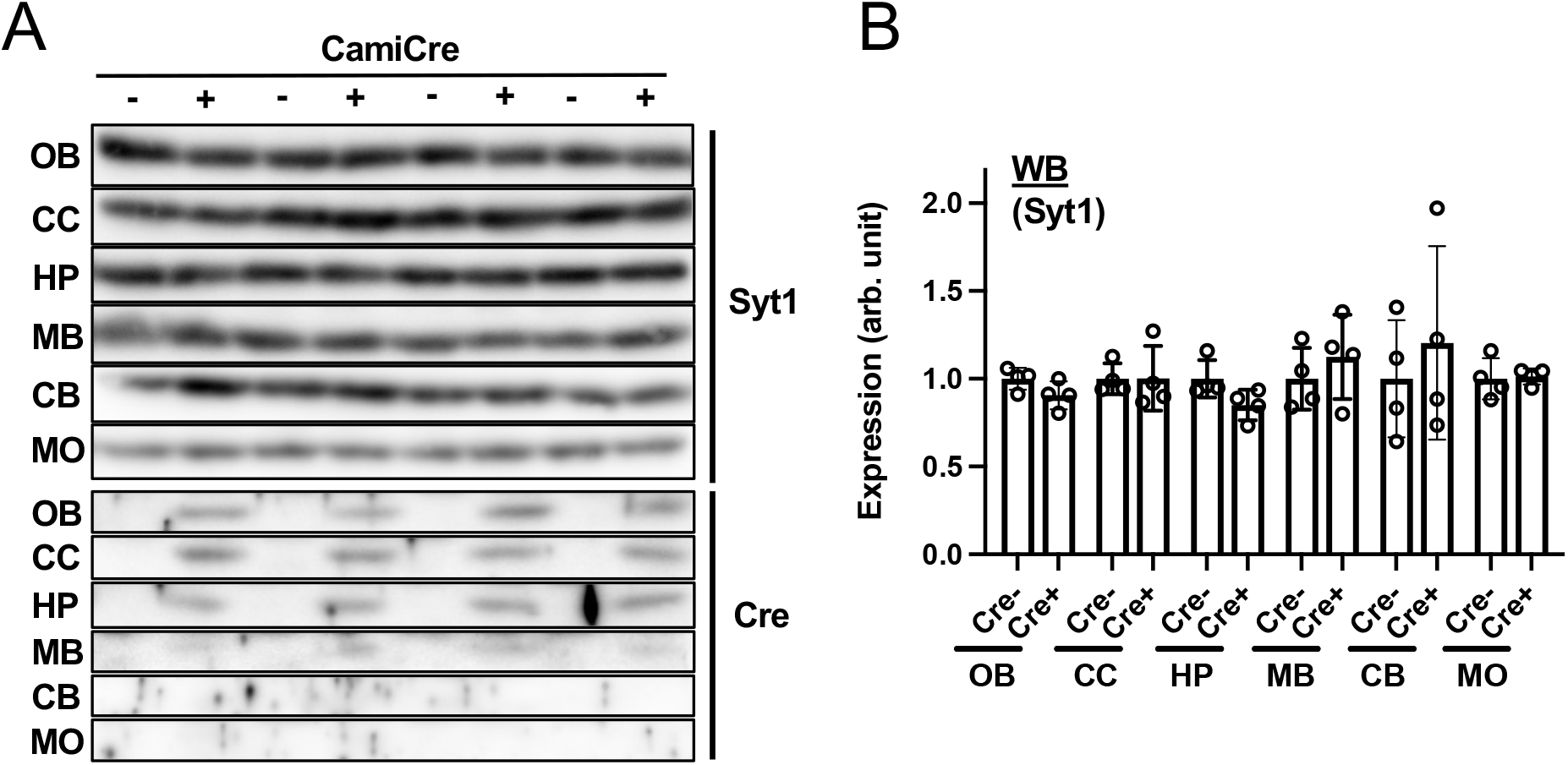
Syt1 expression is unchanged in the CamiCre^+^ brain. (**A**) Western blotting images of different brain regions. CB, cerebellum; CC, cerebral cortex; HP, hippocampus; MB, Midbrain; MO, medula oblongata; OB, olfactory bulb. Cre expression was not detectable in CB and MO. (**B**) Quantification of results in A. Values were normalized by α-tubulin expression level and are shown as fold increases against Cre-controls. Mean ± SD is shown. N = 4. Statistical tests were performed using two-tailed, unpaired t-tests.

**Figure S3.**
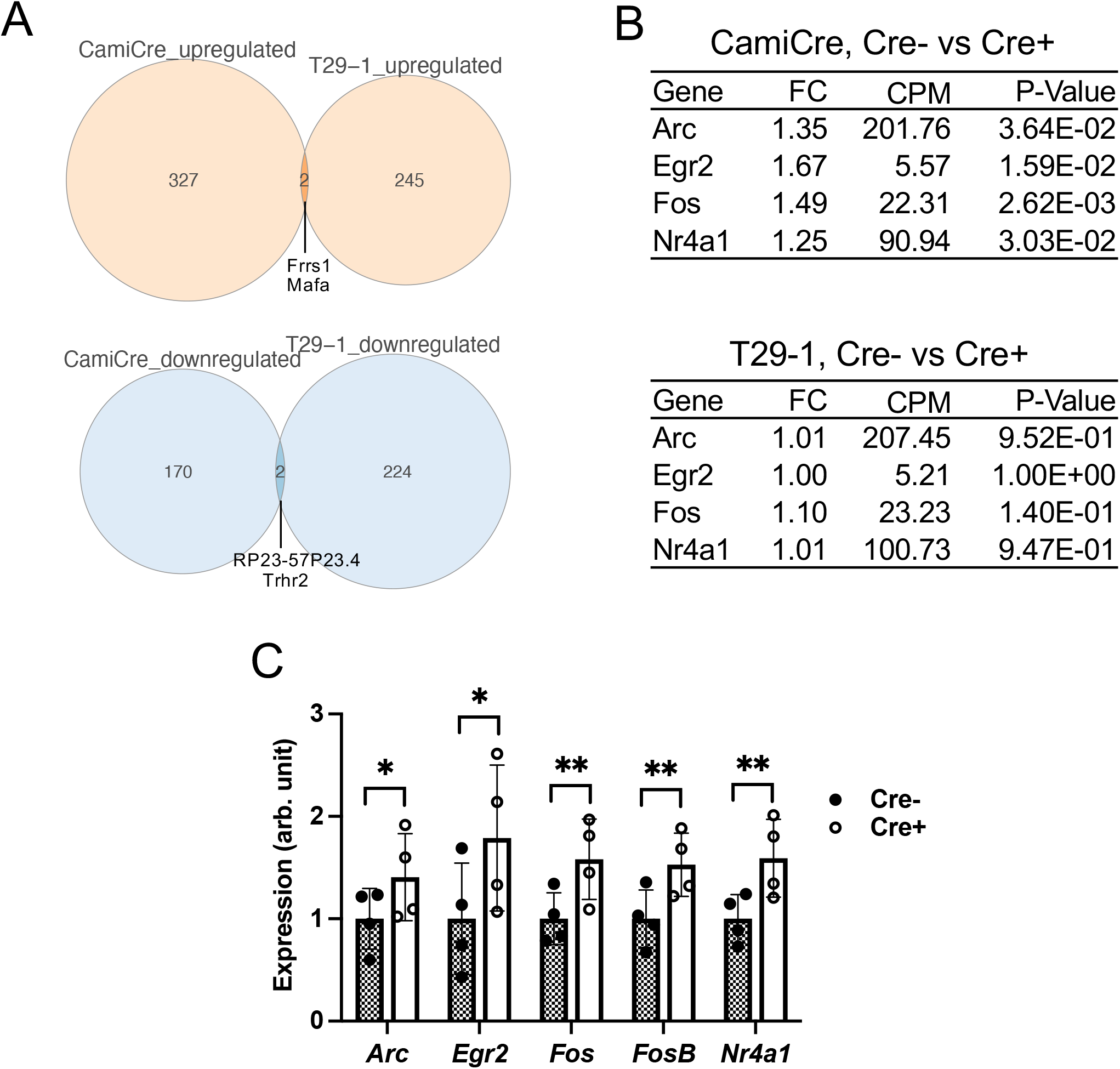
Immediate early genes (IEGs) are upregulated in CamiCre^+^ hippocampi. (**A**) Overlaps of candidate up- or down regulated genes from DE analyses of RNA-seq data. Names of overlapped genes are indicated. (**B**) Differential expression of representative IEGs in CamiCre^+^ hippocampi are shown in the top panel. Results of the same gene set from T29-1 Cre^+^ hippocampi are shown in the bottom panel. N = 4. FC, fold changes of expression in Cre^+^ mice compared to WT littermates; CPM, average counts per millions of all samples; P-value, calculated by exact test. (**C**) qPCR analysis of indicated IEG expression in hippocampi of CamiCre mice. Values were normalized by *Gapdh* expression level and are shown as fold increases against Cre^−^ controls. N = 4. Mean ± SD is shown. *P < 0.05; **P < 0.01. Statistical tests were performed using two-tailed paired t-tests. Littermates raised in the same environment were paired.

**Figure S4.**
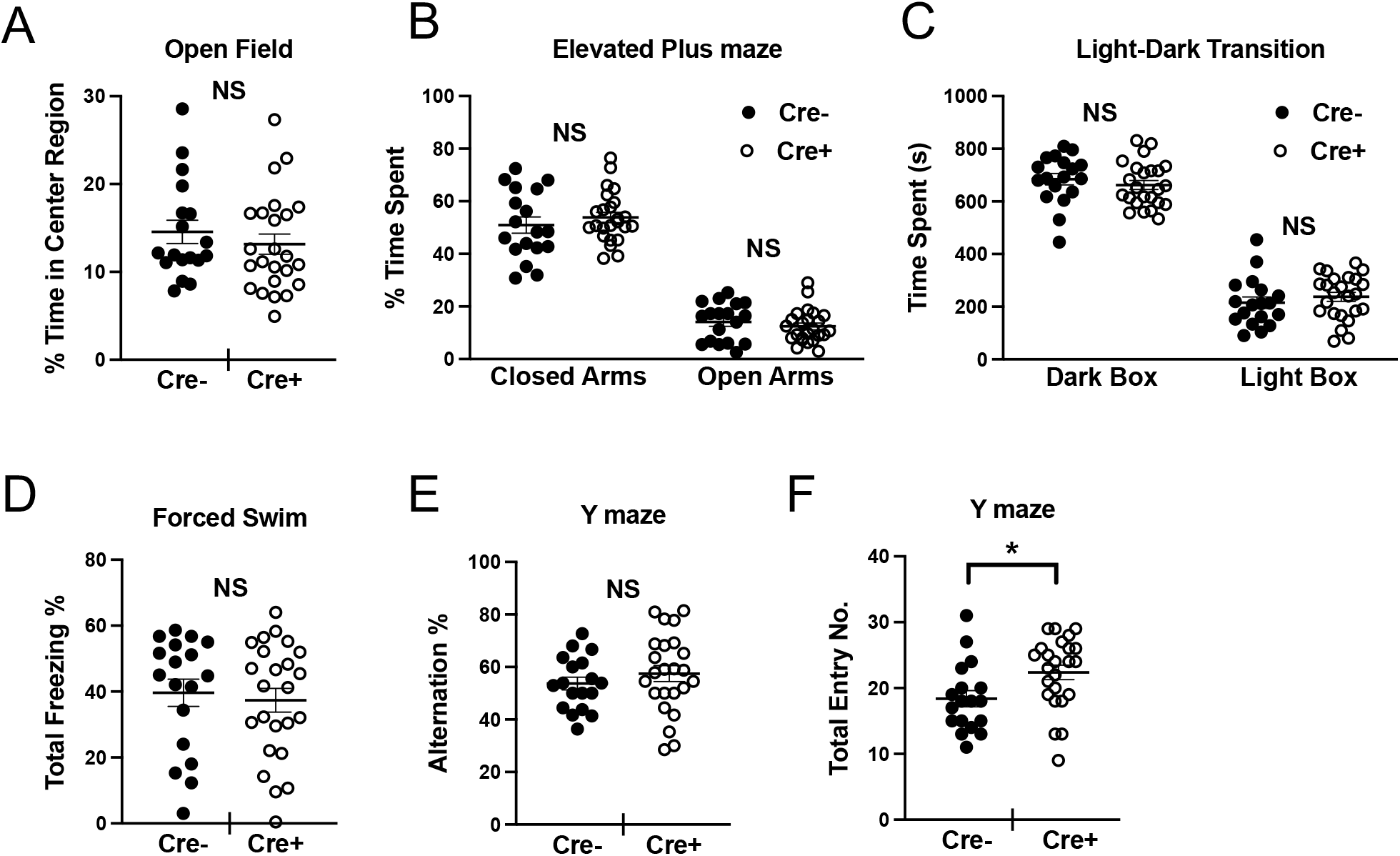
CamiCre^+^ mice show normal anxiety, depression, and short-term working memory. (**A**) Open field test. (**B**) Elevated plus maze. (**C**) Light-dark transition test. (**D**) Forced swim test. (**E**, **F**) Y-maze. Mean ± SEM is shown. NS, not significant; *P < 0.05. N = 18 for Cre^−^, N = 24 for Cre^+^.

## Methods

### Mice

The CamiCre line (B6.FVB-Tg(Camk2a-cre)2Gsc/Cnrm) was obtained from EMMA (EM:01153) and backcrossed seven times onto a C57BL/6J genetic background. The T29-1 line (B6.Cg-Tg(Camk2a-cre)T29-1Stl/J) was obtained from The Jackson Laboratory (JAX: 005359). The Emx1Cre line (B6.129P2-Emx1<tm1(cre)Ito>/ItoRbrc) was provided by Drs. Shigeyoshi Itohara and Takuji Iwasato (RBRC00808). iCre or Cre transgene was kept as heterozygote in all lines. Genotyping was performed by genomic PCR. Primers used were: iCre-FW 5’- gagggactacctcctgtacc −3’, iCre-RV 5’- tgcccagagtcatccttggc −3’; Cre-FW 5’-tcgatgcaacgagtgatgag −3’, Cre-RV 5’- ttcggctatacgtaacaggg −3’, and IL2-FW 5’- ctaggccacagaattgaaagatct −3’; IL2-RV 5’- gtaggtggaaattctagcatcatcc −3’. iCre or Cre primer pair was used in conjunction with IL2 control pair in the ratio of 3:1 concentration. Mice were kept in a 12-h light/12-h dark cycle in a temperature- and humidity-controlled specific pathogen-free vivarium, and they had *ad libitum* access to food and water. 7- to 9-week-old male mice were used for all the experiments. iCre/+ or Cre/+ heterozygous mice and littermate control +/+ mice were used. All animal experiments were conducted according to guidelines for care and use of animals, approved by the Animal Experiment Committee of Okinawa Institute of Science and Technology Graduate University (Approval no. 2016-159 and 2020-283).

### Literature survey

Information was from Mouse Genome Informatics (MGI) database (Bult et al., 2019). We successfully had access to all the links to full text from: http://www.informatics.jax.org/reference/allele/MGI:2181426?typeFilter=Literature. Each paper was surveyed for the description of control mice and those clearly stated the use of Cre^+^ controls were counted.

### qPCR

For qPCR, total RNA was extracted from mouse hippocampi using Isogen II reagent (Nippon Gene). Reverse transcription was performed using PrimeScript II 1st strand cDNA Synthesis Kit (TAKARA) following the manufacture’s protocol. Real-time PCR was performed using TB Green Premix Ex Taq II (Tli RNaseH Plus) (TAKARA) and ViiA7 Real-Time PCR system, Software v1.2 (Applied Biosystems) according to the manufacture’s protocol. For detecting intron in premRNA of Syt2, RT (−) controls were used and confirmed for no detection or detected in higher Ct values with distinct melting curves compared to RT (+) samples. Primers used were: *Gapdh*-FW 5’- ctgcaccaccaactgcttag −3’, *Gapdh*-RV 5’- gtcttctgggtggcagtgat −3’; *Pgk1*-FW 5’- tacctgctggctggatgg −3’, *Pgk1*-RV 5’- cacagcctcggcatatttc −3’; *Syt1*-FW 5’- cagaagacattgcccttgc −3’, *Syt1*-RV 5’- ttttggaatctcctttcttcaatc −3’; *Syt2-Int8*-FW 5’- attggagttggactggaagc −3’, *Syt2-Int8*-RV 5’- cctctgccactgtgcactac −3’; *Syt2*-FW 5’- gacctacaaggcggagagaa −3’, *Syt2*-RV 5’- agcgcaaggaggtacagatg −3’; *Arc*-FW 5’- aggggctgagtcctcaca −3’, *Arc*-RV 5’- tctcagcagccttgagacc −3’; *Egr2*-FW 5’- ttgaccagatgaacggagtgg −3’, *Egr2*-RV 5’- atccaggggtctcttctctcc −3’; *Fos*-FW 5’- agggagctgacagatacactcc −3’, *Fos*-RV 5’- tgcaacgcagacttctcatc −3’; *FosB*-FW 5’- atcgacttcaggcggaaa −3’, *FosB*-RV 5’- ctccaggcgttccttctct −3’; and *Nr4a1*-FW 5’- agcttgggtgttgatgttcc −3’, *Nr4a1*-RV 5’- atgcgattctgcagctcttc −3’.

### Western blotting

Mouse brain regions of interest were quickly dissected on filter paper soaked with ice-cold PBS, snap frozen in liquid nitrogen and stored at −80°C. Brain regions were homogenized in ice-cold lysis buffer (0.3% SDS, 1.67% Triton X-100, 50 mM Tris-HCl pH7.4, 150 mM NaCl, 1 mM EDTA, 1 mM EGTA, 1mM PMSF, 1mM Na_3_VO_4_, 25mM NaF, PI cocktail [5 μg/mL aprotinin, chymostatin, leupeptin and pepstatin A], 10% glycerol) using a pellet mixer, rotated 60 min at 4°C, and centrifuged for 30 min at 20,400 x g, 4°C. The supernatants were collected and adjusted for protein concentration, mixed with 6x SDS sample buffer, and then heated at 56°C for 10 min. Samples were subjected to 8% SDS-PAGE. Western blotting was performed by standard methods. Briefly, proteins were transferred to PVDF membranes (Immobilon, Millipore) and blocked in TBS containing 5% skim milk and 0.1% Tween-20. For antibody dilution, Can Get Signal Immunoreaction Enhancer Solution (TOYOBO) or blocking solution (for α-tubulin and Cre) was used. A chemiluminescent signal was detected using Immobilon (Millipore) or, for α-tubulin, Western Lightning Plus-ECL (Perkin Elmer) on ImageQuant LAS4000 (FujiFilm) following the manufacture’s protocol. Antibodies used were rabbit polyclonal antibody (RpAb) to Syt1 (1:1000) from Synaptic Systems (105103), mouse monoclonal antibody (MmAb) to Syt2 (1:1000) from Abnova (clone ZPN-1), RpAb to Cre (1:1000) from Cell Signaling (15036S), and MmAb to α-tubulin (1:1000) from Sigma-Aldrich (clone DM1A). For reprobing, Restore Plus Western Blot Stripping Buffer (Thermo Scientific) was used. Fiji/ImageJ (NIH) was used for quantification.

### AAV injection

Mice were injected with Cre-expressing adeno-associated virus AAV1.hSyn.Cre.WPRE.hGH (105553-AAV1, Addgene) and GFP-expressing AAV1.hSyn.eGFP.WPRE.bGH (105539-AAV1, Addgene). Stereotaxic surgical procedures were performed as described (Augustinaite and Kuhn, 2020). Viral solutions of around 250 nL were slowly injected using sharp tip beveled quartz pipette into the CA2 region of the hippocampus according to the following coordinates (1.6 mm posterior, 1.6 mm lateral and 1.6 mm ventral to bregma). Additionally, AAV-Cre serial dilutions were done using sterile 0.9% saline solution.

### Immunohistochemistry

Mice were deeply anesthetized with isoflurane and were intracardially perfused with ice-cold sodium phosphate buffer (pH 7.3, NPB), followed by ice-cold 4% paraformaldehyde (PFA) /NPB. The whole brain was removed, separated bilaterally at the medial line and fixed in ice-cold 4% PFA/NPB for 2 h. The brain was further infiltrated sequentially with 10, 15, and 25% sucrose/NPB for more than 4 h at each concentration and then frozen in a Tissue-Tek OCT compound (Sakura Finetek). 10-µm cryosections were attached to an MAS-coated slide glass (S9441 Matsunami) and air dried for 2 h. For permeabilization, sections were incubated in 0.3% Triton X-100/Tris-buffered saline (pH7.5, TBS) for 10 min at room temperature. Sections were blocked with TBS containing a 0.5% blocking reagent (Roche), 2% fetal bovine serum, Mouse Ig Blocking Reagent (Vector Laboratories), and 0.1% Tween-20 for 1 h. Then they were incubated overnight at 4°C with primary antibodies diluted in a dilution buffer (0.5% blocking reagent (Roche), 2% fetal bovine serum, M.O.M. Protein Concentrate (Vector Laboratories), 0.1% Tween-20, TBS). Following washes in TBS containing 0.1% Tween-20 (TBST), sections were incubated for 1 h at RT with secondary antibodies diluted in the dilution buffer. Sections were subsequently stained with DAPI, washed, and coverslipped with Vectashield mounting medium (Vector Laboratories). Sections from CamiCre^+^ and control mice were processed simultaneously on the same slide glass. Antibodies used were RpAb to VGluT1 (1:100) from Millipore (ABN1647), MmAb to Syt2 (1:100) from Abnova (clone ZPN-1), Alexa Fluor 488 Goat Anti-mouse IgG (1:300) and Alexa Fluor 568 Goat Anti-rabbit IgG (1:300) from Invitrogen. Digital images were obtained using Leica TCS SP8 LIGHTNING confocal microscope equipped with motorized stage and LAS X software. Briefly, for low magnification images, the hippocampal region was scanned using HC PL APO CS 40x/0.85 DRY objective and images were stitched automatically using LAS X software. For super-resolution imaging, HC PL APO CS2 63x/1.40 OIL objective was used in LIGHTNING mode with 120 nm resolution. Images with 4928 x 4928 pixels (29 x 29 nm^2^/pixel) were acquired. Original images adjusted only for brightness and contrast by Fiji/ImageJ (NIH) are shown in the figures.

### Colocalization analysis

Colocalization analysis was performed using GDSC ImageJ plug-in according to developer’s Colocalization User Manual (http://www.sussex.ac.uk/gdsc/intranet/microscopy/UserSupport/AnalysisProtocol/imagej/colocalisation). Briefly, super-resolution images stained for Syt2 and VgluT1 in the neuropil region of hippocampal CA1 or CA3 area were acquired as described above. The images (144.72 x 144.72 μm^2^/image) were processed to define foreground and background by Otsu method (or Triangle method for images with relatively low signal-to-noise ratio) using Stack Threshold Plugin. The processed images and original images were used to calculate statistical significance of Manders coefficient and Pearson correlation coefficient, respectively, by Confined Displacement Algorithm Plugin. The pyramidal cell body areas were excluded from the confined region. The significance of correlation or colocalization were calculated by comparing original images to random displacement images. The random displacement was defined using radial displacement chart of Pearson correlation coefficient for each sample according to the user manual. A *P* value of < 0.001 was adopted for statistically significance.

### RNA-seq and DE analysis

Total RNA was purified from mouse hippocampi using Isogen II reagent (Nippon Gene). Subsequently, 500 ng of the total RNA samples were enriched for mRNA using NEBNext Poly(A) mRNA Magnetic Isolation Module and libraries were prepared using NEBNext Ultra II Directional RNA Library Prep Kit for Illumina according to the manufacture’s protocol. All libraries were normalized and pooled for paired-end 150bp sequencing with a single lane of SP flowcell of Illumina NovaSeq 6000. Fastq files containing sequencing reads generated from paired-end RNA sequencing were analyzed using nf-core/rnaseq pipeline version 2.0 (Ewels et al., 2020) to determine read counts using featureCounts (Liao et al., 2014), which were mapped to GRCm38 genome database using STAR aligner (ver. 2.6.1d) (Dobin et al., 2013). The reads were then further analyzed using OmicsBox software (version 1.4.11) for DE analysis using the package EdgeR (ver. 3.11) (Robinson et al., 2010). Reads were normalized using Trimmed Mean of M-values (TMM) normalization method and a cut-off of at least 0.2 count per million (CPM) in two samples was selected. Differentially expressed genes (DEGs) with a *P* value of < 0.05 in exact test, which is based on the quantile-adjusted conditional maximum likelihood (qCML) methods, were used for further analysis. Pathway analyses were performed using Qiagen Ingenuity Pathway Analysis software (ver. 01-19-02).

### Behavioral analysis

Male CamiCre^+^ and WT mice were housed together, with two to five littermates (or mice with close birthdays) per cage after weaning. Mice of 7-8-week-old were acclimated to handling and the experimental room for at least three days before the start of an experiment. Mice underwent a battery of behavioral tests in the following order with at least one day between each test: open field, elevated plus maze, y-maze, light-dark transition, cued and contextual fear conditioning, and forced swim. Experimenters were blinded to the genotype during testing. All experiments were analyzed using an automated system from O’hara & Co. All software for analysis, which are based on the public domain ImageJ program (https://imagej.nih.gov/nih-image/), were from O’hara & Co..

### Open field test

Each subject was placed in the center of an open-field apparatus (50 x 50 x 33.3 cm; W x D x H) illuminated at 100 lux and allowed to move freely for 15 min. Distance travelled in the arena, trace of the movement, and time spent in the center were recorded and analyzed using Time OFCR software.

### Elevated plus maze

The elevated plus maze (EP-3002) consists of two open arms and two closed arms (25 L x 5 W cm) extending from a central area (5 × 5 cm) and is elevated 50 cm from the ground. Mice were placed in the central area of the maze facing one of the open arms under illumination of 100 lux and allowed to move freely for 10 min. The number of entries into open or closed arms and the time spent in each arm were recording and analyzed using Time EPC software.

### Y-maze

The mice were placed at the end of one arm of a Y-shaped maze (YM-3002) with three matte gray PVC arms (3 x 40 x 12 cm; W x D x H) at a 120° angle from each other, illuminated at 50 lux for 10 minutes. Entry sequence into each arm and the total number of arm entries were recorded using Time YM software. An alternation is defined as the entry into all three arms on consecutive choices. An entry occurs when all four limbs of the mouse are within the arm. The alternation score (%) was calculated using the following 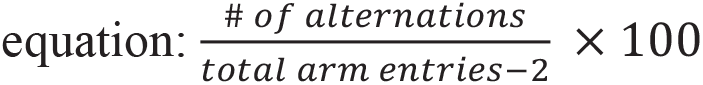.

### Light-dark transition test

The apparatus consisted of a box (20 x 20 x 25 cm) divided into two compartments of equal size by a partition with a door: a white polyvinylchloride chamber illuminated at 500 lux and a black polyvinylchloride chamber without illumination. Mice were placed in the dark compartment and allowed to move freely with the door open for 15 min. The total number of transitions, time spent in each side, and latency to enter the light side were recorded and analyzed automatically using Time LD software.

### Contextual and cued fear conditioning

Fear conditioning was conducted in a conditioning chamber (10 x 10 x 10 cm) surrounded by a sound-attenuating chamber. On day 1, mice were placed in the conditioning chamber (CL-3002L) for 120 s as habituation and then presented with 3 tone-shock pairs at 90 seconds intervals. The tone-shock pair consists of a tone (65-dB/10-kHz) for 30 seconds and a foot shock of 0.5mA during the last 2 s of the tone. Following the last tone-shock pair, the mice were left in the chamber for 90 s. On day 2, mice were placed in the conditioning chamber without any tone-shock pairs for 360 s to test contextual learning. On day 3, the mice were placed in a novel chamber (CLT-3002L) for 180 s and then presented with a tone for 180 s to test cued learning. Freezing responses were automatically recorded during each test and analyzed using Image FZC 2.22 sr2 software.

### Forced swim test

Mice were placed in a Plexiglas cylinder (20 cm H x 11.4 cm D) filled with water (23-25°C) under 300 lux illumination. Duration of immobility (freezing) was recorded automatically during a 10 min test session using the ImagePS/TS software. Percentage of immobility was calculated for the last 8 minutes. After testing, mice were removed and dried before being placed in their home cage.

### Statistical analysis

Statistical analyses in this work employed unpaired two-tailed Student’s *t* tests, two-tailed Welch’s *t* tests, two-tailed paired *t* tests, or a two-way ANOVA with Geisser-Greenhouse correction, where appropriate using GraphPad Prism 9. A *P* value of < 0.05 was considered statistically significant.

